# Manipulating the rapid consolidation periods in a learning task affects general skills more than statistical learning

**DOI:** 10.1101/2022.05.05.490763

**Authors:** Laura Szücs-Bencze, Lison Fanuel, Nikoletta Szabó, Romain Quentin, Dezso Nemeth, Teodóra Vékony

## Abstract

Memory consolidation processes have traditionally been investigated from the perspective of hours or days. However, the latest developments in memory research showed that memory consolidation processes could occur even within seconds, possibly due to the neural replay of just-practiced memory traces during short breaks. Here, we investigate this rapid form of consolidation during statistical learning. We aim to answer (a) whether this rapid consolidation occurs in implicit statistical learning and general skill learning and (b) whether the duration of rest periods affects these two learning types differently. Participants performed a widely used statistical learning task - the Alternating Serial Reaction Time (ASRT) task - that enables us to measure implicit statistical and general skill learning separately. The ASRT task consisted of 25 learning blocks with a rest period between the blocks. In a between-subjects design, the length of the rest periods was fixed at 15 or 30 seconds, or the participants could control the length themselves. We found that the duration of rest periods does not affect the amount of statistical knowledge acquired but does change the dynamics of learning. Shorter rest periods led to better learning during the learning blocks, whereas longer rest periods promoted learning also in the between block rest periods, possibly due to the higher amount of replay. Moreover, we found weaker general skill learning in the self-paced group than in the fixed rest period groups. These results suggest that distinct learning processes are differently affected by the duration of short rest periods.

## Introduction

Learning is the process of gaining knowledge or skills by studying, practicing, or experiencing events repeatedly. The development of knowledge is not limited to the duration of the practice, as it continues to develop between training sessions, either during awake or sleep periods. This phenomenon is known as memory consolidation (Robertson et al., 2004a). Consolidation was previously thought to occur during an extended period, from hours to days (Squire et al., 2015). Recent studies suggest that memory consolidation can occur within shorter periods, even in seconds (Bönstrup et al., 2019). It has been suggested that this phenomenon is due to the neural replay of just-practiced memory traces during short breaks (Buch et al., 2021). However, research to date has investigated the effect of short rest periods – when replays occur – on learning only with one predetermined fixed rest period (Bönstrup et al., 2019; Du et al., 2016) or multiple self-paced rest periods (Fanuel et al., 2022; Quentin et al., 2021). Their results do not allow us to determine the causal role of short rests in the learning process, nor whether more replay leads to better learning performance. To fill this gap, in the present study, we manipulated the duration of rest periods – indirectly the possible amount of replay – to test whether rest durations affect differently (1) the general speed-up on a statistical learning task independently of the statistical probabilities in the task (general skill learning) (2) and the learning of statistical probabilities (statistical learning).

Rest periods inserted in a learning process may facilitate the acquisition of new skills (Walker et al., 2002). In the study of Bönstrup and colleagues (2019), the performance on an explicit motor-skill learning task improved during short, 10-second rest periods. In their study, frontoparietal beta oscillatory activity during rest periods was associated with learning gains from rapid consolidation. A re-analysis of these data suggested that such rapid consolidation is driven by the replay of just-practiced memory traces during short breaks (Buch et al., 2021). As the awareness of learning determines how knowledge acquisition gains from offline periods (Robertson et al., 2004a, 2004b), this rapid consolidation may manifest differently or even be absent during implicit learning of statistical regularities. Implicit statistical learning implies the process of unintentional acquisition of probabilistic regularities embedded in the environment (Cleeremans and Jimenez, 1998; Howard et al., 2004). So far, two studies have examined the rapid consolidation in implicit statistical learning. On one hand, they have found that implicit acquisition of statistical knowledge does not improve during short breaks but deteriorates, and this effect is not associated with the length of rest periods (Fanuel et al., 2022). On the other hand, statistical learning was found to develop during practice (online) indicating that it benefits from evidence accumulation during practice and the information learned does not consolidate during short rest periods (Quentin et al., 2021).

Based on previous results, implicit statistical learning occurs only online and does not benefit from rapid consolidation. However, previous studies have a crucial limitation: the length of the rest periods was not controlled experimentally. In the present study, to grasp a causal relationship between the length of breaks and the learning performance, we varied the rest period duration between participants. We aimed to test whether shorter and longer rest durations affect the performance of implicit statistical learning (i.e., the learning of probabilistic regularities) and general skill learning (i.e., the general speed-up on a learning task independently of the statistical probabilities). To tackle this question, we used the Alternating Serial Reaction Time (ASRT) task (Howard et al., 2004), which enables us to measure these two aspects of learning separately. Healthy adults performed 25 blocks of the ASRT task (one block = 80 trials) and were offered to rest after each block. The rest period was either (1) a shorter 15-second break, (2) a 30-second break, or (3) a self-paced duration. As rapid consolidation is related to neural replay (Buch et al., 2021), we expected that the extended rest period will benefit implicit statistical learning more than the shorter rest period, due to multiple of replays. Moreover, we aimed to test whether there is dissociation in the temporal dynamics of general skill learning and statistical learning regarding online and offline changes and how it is affected by the length of the rest periods.

## Methods

### Participants

Three hundred and sixty-one participants participated in this preregistered, online study (https://osf.io/pfy7r). Participants were university students and gained course credits for their participation. Following careful quality control of participant data (see below in section *Quality control of data*), the final sample consisted of 268 participants (*M_age_* = 21.46 years, *SD_age_* = 2.20 years; 77.61% female, 22.39% male). Participants were randomly divided into three groups (15-second break, 30-second break, and self-paced). Participants of the three groups did not differ in age, education, sex, handedness, or working memory performance (see Table 1). All participants had normal or corrected-to-normal vision, and none of them reported a history of any neurological or psychiatric condition. Participants provided informed consent, and all study procedures were approved by the Research Ethics Committee of the Eötvös Loránd University, Budapest, Hungary, and was conducted in accordance with the Declaration of Helsinki.

**Table 1.** Descriptive statistics of the three experimental groups

|  | **Self-paced**<br>**Group (n = 88)** | **15-sec Group**<br>**(n = 90)** | **30-sec Group**<br>**(n = 90)** | **Comparison** |
| --- | --- | --- | --- | --- |
| **Age (years)** | 21.89 ± 2.07 | 21.24 ± 2.23 | 21.27 ± 2.26 | $p = .09, BF_{01} = 2.76$ |
| **Education (sec/BA/MA)** | 68/17/3 | 67/22/1 | 73/16/1 | $p = .56, BF_{01} = 16.37$ |
| **Gender (m/f)** | 18/70 | 20/70 | 22/68 | $p = .82, BF_{01} = 16.43$ |
| **Handedness (l/r/a)** | 8/80/0 | 8/80/2 | 10/78/2 | $p = .68, BF_{01} = 10.02$ |
| **2-back (d-prime)** | 1.57 ± 0.95 | 1.49 ± 0.87 | 1.52 ± 0.89 | $p = .82, BF_{01} = 21.03$ |

Means and standard deviations for age, and 2-back task are presented. For education (sec = secondary education or lower, BA = bachelor's level or equivalent, MA = master's level or equivalent), gender (m = male, f = female) and handedness (l = left, r = right, a = ambidextrous) case numbers are presented. To compare age and 2-back task scores, one-way ANOVAs were conducted. To compare education, gender, and handedness ratios, Chi-square tests were conducted.

### Alternating Serial Reaction Time (ASRT) task

We used the ASRT task to measure implicit statistical and general skill learning separately. The ASRT task was programmed in JavaScript with the jsPsych framework (de Leeuw, 2015) (code is openly available on GitHub, https://github.com/vekteo/ASRT_rapid_consolidation) (Vékony, 2021). In the ASRT task, a visual stimulus appeared on the screen (a drawing of a dog’s head) in one of four horizontal locations. Participants must have indicated the location of the target stimulus by pressing the corresponding key on the keyboard (from left to right, the S, F, J, and L keys on a computer keyboard). Participants were instructed to use their left and right middle and index fingers to respond to the targets. Unknown to the participants, the stimuli followed a probabilistic eight-element sequence, where pattern and random elements alternated with each other (e.g., 2 – r – 1 – r – 3 – r – 4 – r, where *r* indicates a random location, and the numbers represent the predetermined positions from left to right). Each participant was randomly assigned to one of 24 possible sequences (as the permutation of the 8-element sequence structure allowed 24 different sequences) and then was exposed to that one sequence throughout the task. Due to the probabilistic sequence structure, some runs of three consecutive stimuli (triplet) appeared with higher probability (high-probability triplets) because the third element of a triplet can be predicted by the first trial with a greater probability (62.5 % of all trials) compared to other third elements that can be predicted by the first trial with a lower probability (low-probability triplets, 37.5 % of all trials). We can assort every item according to whether they are the third element of a high- or a low-probability triplet. Statistical learning was defined as the increase of reaction time difference between trials that were the third element of a high-probability triplet or low-probability triplet. General skill learning was defined as the overall speeding up on the task (i.e., smaller RTs in later blocks), regardless of the probability of occurrence of items.

### Procedure

We used the Gorilla Experiment Builder (https://www.gorilla.sc) to host our experiment (Anwyl-Irvine et al., 2020), which allows accurate stimulus and response timing in online experiments (Anwyl-Irvine et al., 2021). Data was collected between 13 April 2021 and 31 October 2021 (experiment material is available on Gorilla Open Materials, https://app.gorilla.sc/openmaterials/397611). Participants were randomly assigned to one of three versions of the task, which differed only in the duration of between-block rest periods. The between-block rest periods were either (1) 15-second breaks, (2) 30-second breaks, or (3) self-paced (i.e., participants were allowed to continue the task with the next block whenever they were ready). The participants performed two practice blocks, then continued with 25 learning blocks, which took approximately 25 min to complete. Each block consisted of 80 trials, corresponding to the eight-element sequence repeated ten times. Accuracy and reaction time (RT) were recorded for each trial. After accomplishing the ASRT task, we tested the participants’ awareness of the hidden structure with a short questionnaire and a task based on the Process Dissocation Procedures, which enables us to differentiate explicit and implicit processes in memory tasks (Jacoby, 1991). The results revealed that the knowledge of the statistical regularities remained equally implicit for all groups (see details in the “Process Dissociation Procedures Task” section of the Supplementary Materials) (Vékony, 2021). Finally, they performed 0-back and 2-back tasks (Kirchner, 1958) to assess their working memory capacity (Vékony, 2021). Data are available on OSF (https://osf.io/ukbfz/).

### Quality control of data

We have set up exclusion criteria before the analysis of the data. Participants were deemed unreliable and were excluded if 1) they did not reach 80% accuracy on the ASRT task (34 participants were excluded for this reason), as in lab experiments, the general accuracy on the ASRT task is typically over 90% (Janacsek et al., 2012), or 2) perform the 0-back task with less than 60% accuracy (8 participants), 3) did not complete the n-back tasks correctly (i.e., did not press response keys during the task) (16 participants), or 4) had quitted the experiment and restarted later (4 participants), or 5) indicated that had already taken part in an ASRT experiment (8 participants), and 6) had not started blocks on time after the rest period expired (21 participants). We fixed this limit at 1500 ms after the end of the rest period in at least five blocks, and we also excluded the participants whose average RT for the blocks’ first trials was above 1000 ms (9 participants in the 15-sec group and 12 participants in the 30-sec group). Besides the participants excluded according to the pre-determined exclusion criteria, as the age range was wide and unequal between the groups, outlying participants (age > 35 years) were also excluded (11 participants). To ensure that the non-preregistered exclusion criterion did not affect our results, all analyses were also conducted with these participants; these analyses led to similar results (see details in the “Results without age-based exclusion” section in Supplementary Materials).

### Quantification of statistical learning and general skill

Inaccurate responses, trills (e.g., 1-2-1) and repetitions (e.g., 1-1-1), and trials with an RT of more than 1000 ms, were excluded from the analysis. There was a total of 535.994 trials, from which a total of 49.927 (9.31 %) incorrect trials were excluded. Regarding triplets, 48.715 (9.09 %) was trills (e.g., 1-2-1) and 16.302 (3.04 %) was repetitions (e.g., 1-1-1) of all trials. Furthermore, there were 1304 (0.24 %) of the trials with RT above 1000 ms. As there were overlaps between the trials with different exclusion criterias (e.g., from the 48.715 trills 7544 was also incorrect trials), the total amount of excluded trials is not the sum of the numbers of the different types of exluded trials. We excluded a total amount of 108.344 trials (20.22 % of all trials).

To facilitate data processing and filter out noise, the blocks of ASRT were organized into units of five consecutive blocks (Barnes et al., 2008; Bennett et al., 2007; Nemeth et al., 2010), for which we calculated statistical learning and general skill learning scores. Each task trial was categorized as the third element of a low- or a high-probability triplet (except the first two trials of each block that could not have been categorized as the third element of a high- or low-probability triplet). To measure the degree of implicit statistical learning, we calculated a statistical learning score by subtracting the high-probability triplets’ median RT from the low-probability triplets’ median RT. Then, to control the difference in base RTs between groups, we divided this learning score by the mean RT (standardized statistical learning scores). To measure general skill learning, the median RTs of each unit of five blocks were calculated regardless of the probability of occurrence of items.

### Quantification of online and offline changes

Further scores were calculated to compare the online and offline general skill learning and statistical learning changes. Each block of 80 trials was divided into five bins (each containing 16 consecutive trials). For each bin, we calculated the difference between high vs. low-probability triplets, resulting in a single learning score for each bin for each block. To calculate the online change in statistical learning, we subtracted the learning score of the first bin from that of the last bin of the same block (the change from the beginning to the end of the block). Twenty-five scores were obtained corresponding to the online changes of learning in the 25 blocks. We averaged over the 25 online learning scores to obtain a single online learning score for each participant. To calculate the offline change of statistical learning, we subtracted the learning score of the last bin from the first bin of the next block (the change from the end of the block and the beginning of the next block). Twenty-four scores were obtained corresponding to the offline changes of learning in the 25 blocks (henceforth referred to as *change scores*). We averaged over the 24 offline learning scores to obtain a single offline learning score for each participant. The same procedure was repeated to obtain the online and offline changes for general skill learning, except that scores were obtained from median RTs independently of the probability of items.

### Statistical analysis

Before conducting the statistical analyses of the main hypotheses, we had calculated the mean and median rest duration of the self-paced group. The mean rest period in the self-paced group was 16.67 s (*SD* = 25.48), and the median rest period was 10.58 s. One-sample t-tests revealed that the mean rest duration of the self-paced group did not differ significantly from the rest duration of the 15-sec group (*t*(87) = 0.62, *p* = .54), but significantly differed from the rest duration of the 30-sec group (*t*(87) = −4.91, *p* < .001).

The learning blocks were grouped into five larger units of analysis (Blocks 1-5, Blocks 6-10, Blocks 11-15, Blocks 16-20, and Blocks 21-25). Statistical analysis was performed in JASP 0.16 (JASP Team, 2021). Mixed-design ANOVAs on median RTs and statistical learning scores were performed to compare general skill learning and implicit statistical learning between groups, respectively. Offline and online changes were also compared with mixed-design ANOVAs separately for statistical learning and general skill learning. Greenhouse-Geisser corrections were applied if necessary. Significant main effects and interactions were further analyzed using Bonferroni-corrected post-hoc comparisons.

In addition to the classical frequentist approach, Bayesian ANOVAs were also performed with the same factors as described above. Here, we report the exclusion Bayes factors (BF) of Bayesian Model Averaging across all matched models. *BF*_exclusion_ indicates the amount of evidence for the exclusion of a given factor. Accordingly, the higher the *BF*_exclusion_ value (above 1), the more it supports the exclusion of the given factor, and vice versa, the smaller the *BF*_exclusion_ value (below 1), the more evidence of inclusion.

## Results

### Did rest period duration influence statistical learning?

To test whether the duration of rest periods between learning blocks affected statistical learning, we conducted a mixed-design ANOVA with the within-subjects factor of Blocks (Blocks 1-5 vs. Blocks 6-10 vs. Blocks 11-15 vs. Blocks 16-20 vs. Blocks 21-25) and the between-subjects factor of Group (self-paced, 15-sec breaks, 30-sec breaks) on the learning scores. The analyses revealed a gradual increase of learning scores in each group, irrespective of the rest period duration [main effect of Blocks, *F*(4, 1060) = 25.68, *p* < .001, η_*p*_^2^ = .09, *BF*_exclusion_ < 0.001]. According to pairwise comparisons, there was no significant increase in learning between Blocks 6-10 and Blocks 11-15 (*p* = .82), between Blocks 6-10 and Blocks 16-20 (*p* = .06), between Blocks 11-15 and Blocks 16-20 (*p* < .99), and between Blocks 16-20 and Blocks 21-25 (*p* = .19). All other paired comparisons of block arrays were significant (all *p* < .01). Thus, the consecutive learning units did not significantly differ from each other but learning could be discovered between temporally more distant parts of the task. Importantly, the three experimental groups did not differ in statistical learning [main effect of Group, *F*(2, 265) = 0.65, *p* = .53, η_*p*_^2^ < .01, *BF*_exclusion_ = 31.39]. The Blocks × Group interaction was also non-significant, *F*(8, 1060) = 0.28, *p* = .97, η_*p*_^2^ < .01, *BF*_exclusion_ = 3 262.88, thus the three groups did not differ in the time course of statistical learning either (see Figure 2A and Figure 2C) (for the performance on the original variables see the section, Performance of high- and low-probability triplets in the three groups” in Supplementary Materials).

**Figure 1.**
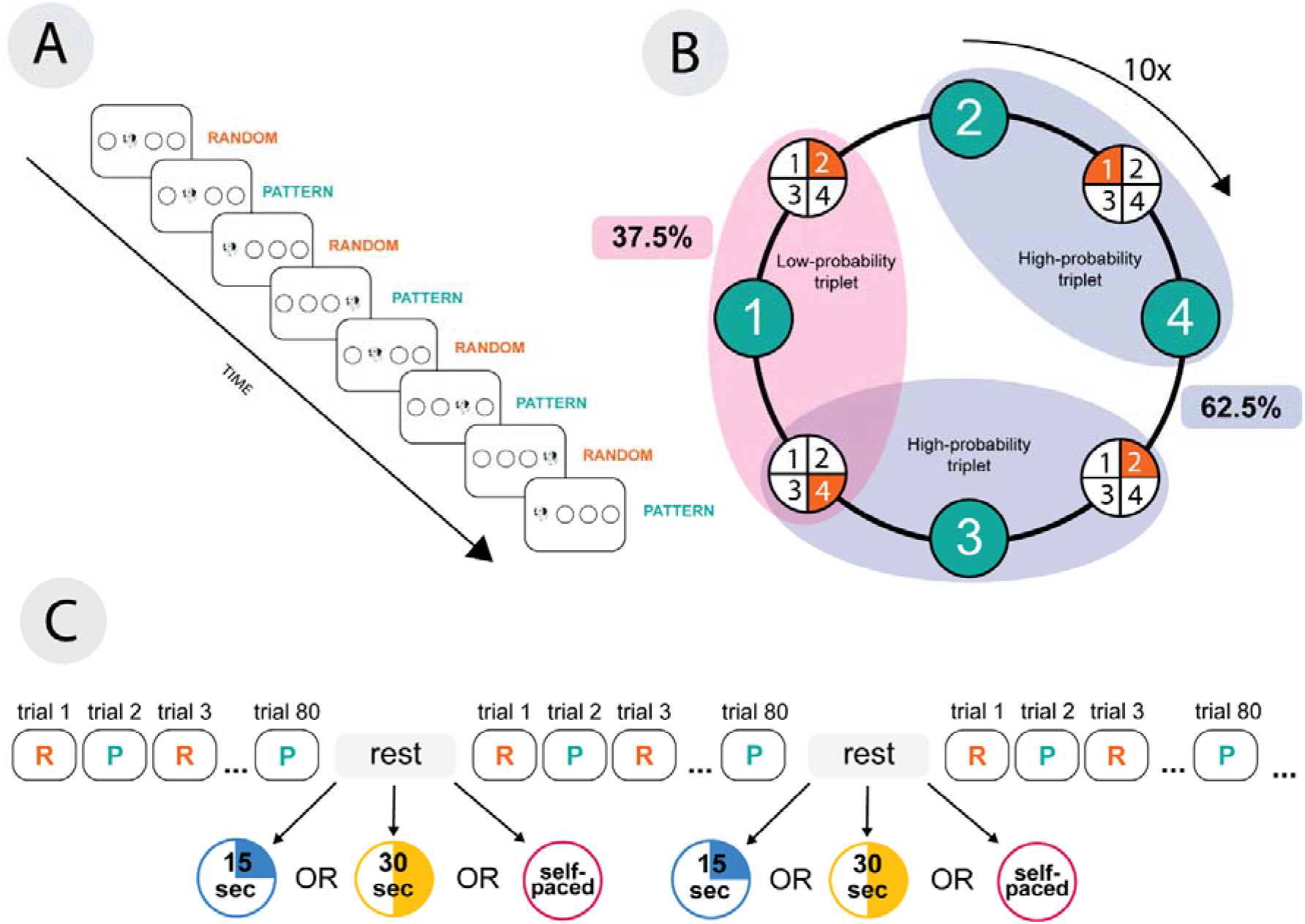
The Alternating Serial Reaction Time (ASRT) task and the study design. (A) The temporal progress of the task. A drawing of a dog’s head appeared as a target stimulus in one of four horizontally arranged locations. The stimuli followed a probabilistic sequence, where every other trial was a part of a four-element fixed sequence (pattern elements) interspersed with random elements. (B) The formation of triplets in the task. In the eight-element probabilistic sequence, pattern (green) and random (orange) trials alternated. Numbers 1-4 represent the location of the four circles from left to right. Every trial was categorized as the third element of three consecutive trials (i.e., a triplet). Due to the probabilistic sequence structure, some triplets appeared with higher probability (high-probability triplets) than others (low-probability triplets). The ratio of high-probability triplets was higher (62.5% of all trials) than that of low-probability triplets (37.5% of all trials). The eightelement alternating sequence was repeated ten times in a learning block. (C) Study design. Each block contained 80 trials. The between-blocks rest period was either 30 seconds (30-sec group), 15 seconds (15-sec group), or a self-paced duration (self-paced group).

**Figure 2.**
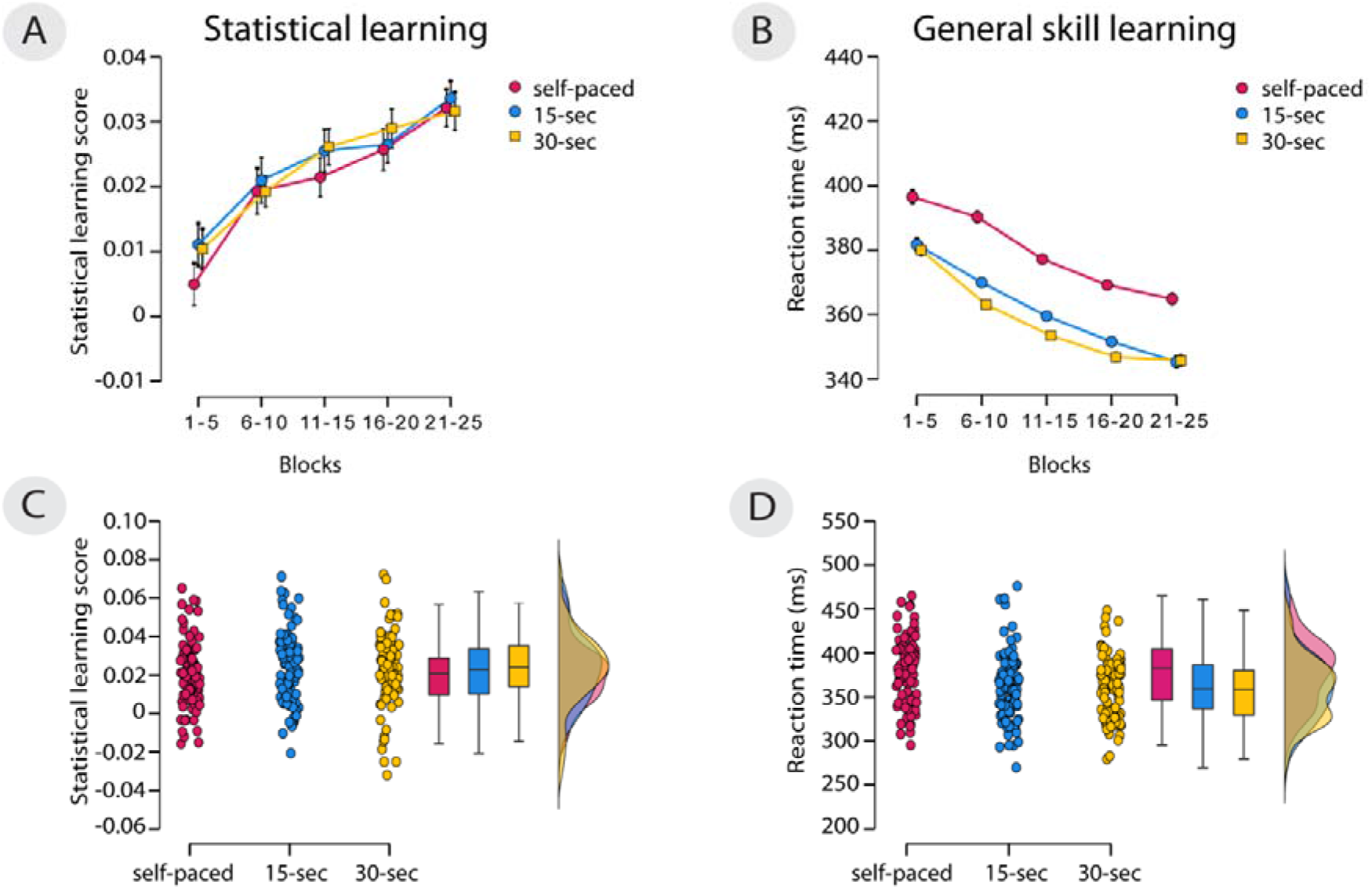
The effect of manipulating rest period duration on statistical learning and general skill learning. Error bars represent the standard error of the mean. The x-axis indicates the blocks of the experiment; the y-axis represents the statistical learning score/reaction time. (A) The temporal dynamics of statistical learning scores in the three groups. All groups showed a significant increase in statistical learning throughout the experiment, but the learning of the three groups did not differ. (B) The temporal dynamics of general skill learning in the three groups. All groups showed a decrease in RT over the course of the experiment, suggesting learning of general skills. Self-paced group showed slower RTs compared to the 15-sec group and 30-sec group. (C) Individual data of the overall statistical learning scores (one dot represent the mean statistical learning score of the epochs for one participant). Boxplots and violin plots visualize the distribution of statistical learning scores in the three groups. (D) Individual data of the general RT scores (one dot represent the mean general RT of the epochs for one participant). Boxplots and violin plots visualize the distribution of general RT in the three groups.

### Did rest period duration influence the performance of general skill learning?

To test whether the overall speed-up on the task differed between groups (i.e., whether the duration of rest periods between learning blocks affected general skill learning), we conducted a mixed-design ANOVA with the within-subjects factor of Blocks (Blocks 1-5 vs. Blocks 6-10 vs. Blocks 11-15 vs. Blocks 16-20 vs. Blocks 21-25) and the between-subjects factor of Group (self-paced, 15-sec breaks, 30-sec breaks) with median RT as the dependent variable. We found a gradual decrease in RTs throughout the task [main effect of Blocks, *F*(2.73, 723.72) = 275.21, *p* < .001, η_*p*_^2^ = .51, *BF*_exclusion_ < 0.001]. Based on pairwise comparisons, every epoch significantly differed from each other (all *p* < .01), with increasing learning through all blocks. The three groups significantly differed in response times [main effect of Group, *F*(2, 265) = 8.69, *p* < .001, η_*p*_^2^ = .06, *BF*_exclusion_ = 0.01], with the self-paced group being slower than the 15-sec and 30-sec groups. The Blocks × Group interaction was also significant, *F*(8, 1060) = 2.33, *p* = .04, η_*p*_^2^ = .02, *BF*_exclusion_ = 5.25]. Pairwise comparisons revealed significantly higher RTs in the self-paced compared to the 30-sec group in Blocks 6-10, Blocks 11-15, Blocks 16-20 and Blocks 21-25 (all *p* < .01). The self-paced group also showed significantly higher RTs compared to the 15-sec group in Blocks 6-10, Blocks 11-15, Blocks 16-20, and Blocks 21-25 (all *p* < .01). Thus, the three groups showed a similar speed in the first learning unit, but the self-paced group began to slow down compared to the other two groups starting from the second learning unit (see Figure 2B and Figure 2D). However, the *BF*_exclusion_ score of the interaction is above 3, which indicates moderate evidence for the lack of interaction; thus, the interaction is deemed unreliable.

### How did break duration affect offline and online statistical learning?

A mixed-design ANOVA was run with the within-subjects factor of Learning Phase (offline vs. online) and the between-subject factor of Group (self-paced, 15-sec breaks, 30-sec breaks) on the change scores of statistical learning. The ANOVA revealed an interaction between Learning Phase and Group factors, *F*(2, 265) = 3.51, *p* = .03, η_*p*_^2^ = .03, *BF*_exclusion_ = 0.05. Bonferroni-corrected post-hoc comparisons revealed that online and offline changes differed in the 15-sec break group (*p* = .04): the offline changes were significantly smaller than the online changes. No main effect of Group [*F*(2, 265) = 1.60, *p* = .20, η_*p*_^2^ = .01, *BF*_exclusion_ = 42.28] or Learning Phase was found [*F*(2, 265) = 2.50, *p* = .12, η_*p*_^2^ < .001, *BF*_exclusion_ = 0.97] (Figure 3) (for dynamic change of offline and online learning scores see the, Offline and online changes in statistical learning across all blocks” section of Supplementary Materials).

**Figure 3.**
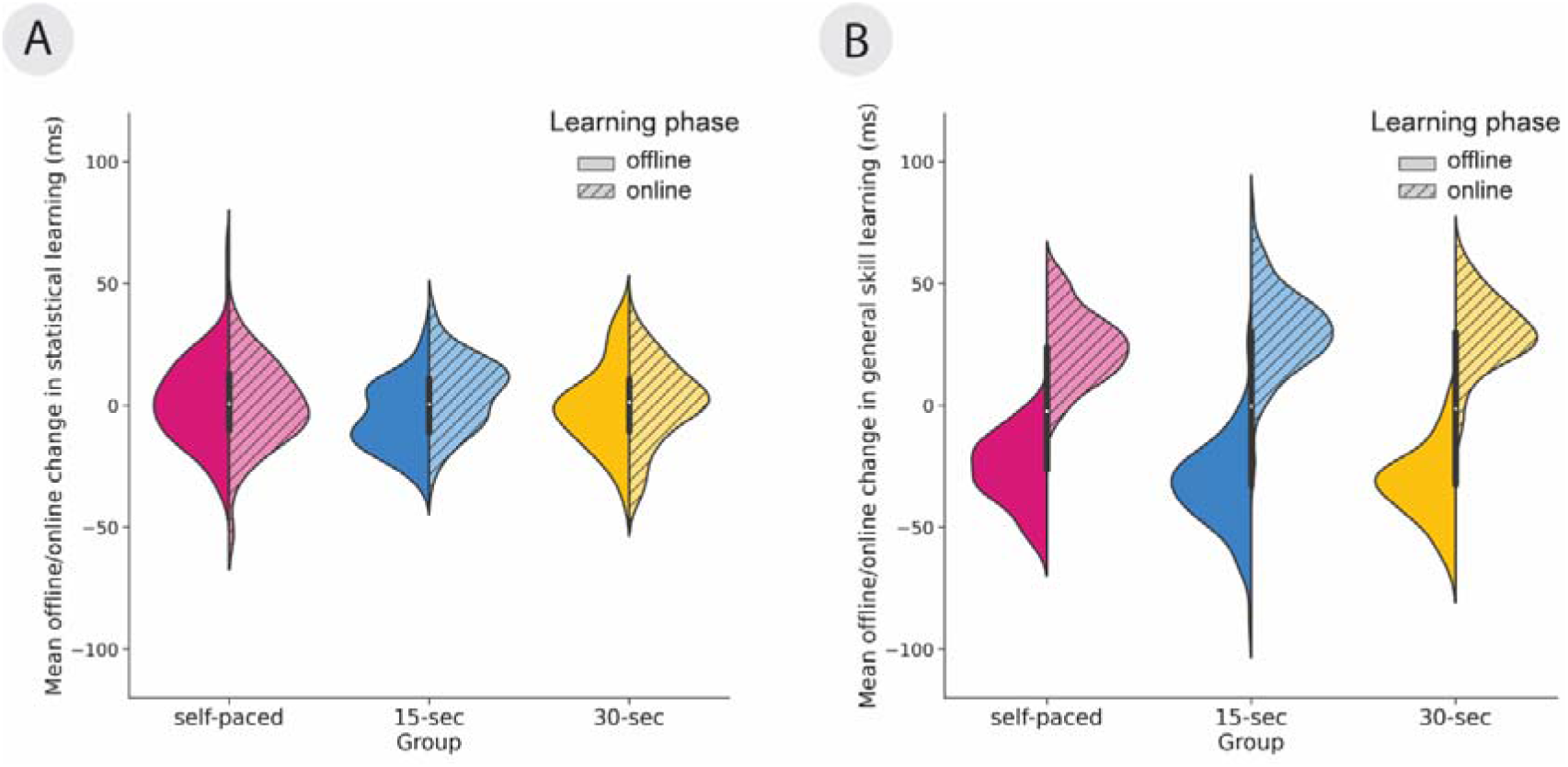
The offline vs. online changes in statistical learning. The x-axis indicates the three groups, and the y-axis represent the mean offline/online changes in ms. The filled halves of the violin plots indicate offline changes, whereas the striped halves the online changes. (A) In the 15-sec group, the offline and online changes differed from each other in statistical learning: the online changes were significantly higher than the offline changes. The group-level online changes were positive (indicating online improvement in statistical learning), whereas the offline changes were negative (indicating forgetting). (B) Changes in general skills were similar in the three groups: acceleration after the offline periods and deceleration during the online period.

To clarify whether offline and online learning occured in the whole sample as well as in the three groups, one sample t-tests were conducted. We have found that on the whole sample, the online learning score was significantly different from zero (*t*(267) = 2.05, *p* < .05), while the offline learning score was not (*t*(267) = 1.11, *p* = .27). In the self-paced group, neither the online learning score (t(87) = −0.17, p = .86, nor the offline learning scores (*t*(87) = 0.92, *p* = .36) differed from zero. Similarly in the 30-sec group, neither the online learning scores (*t*(89) = 0.61, *p* = .55), nor the offline learning score (*t*(89) = −0.05, *p* = .96) differed from zero. However, both learning scores differed from zero in the 15-sec group: the online learning scores were higher than zero (*t*(89) = 3.50, *p* < .001), whilst the offline learning scores were smaller than zero (*t*(89) = −3.39, *p* < .01). We can conclude that in this group, participants learned online and forgot offline. We suggest that the reason behind the lack of online and offline learning at group level in the other two groups can be explained by the balanced ratio of positive and negative learning scores within the groups (see Figure 4A, Figure 4B and Figure 4C) (see details in the, Dynamics of offline/online statistical learning and forgetting in the different groups” section of the Supplementary Materials).

**Figure 4.**
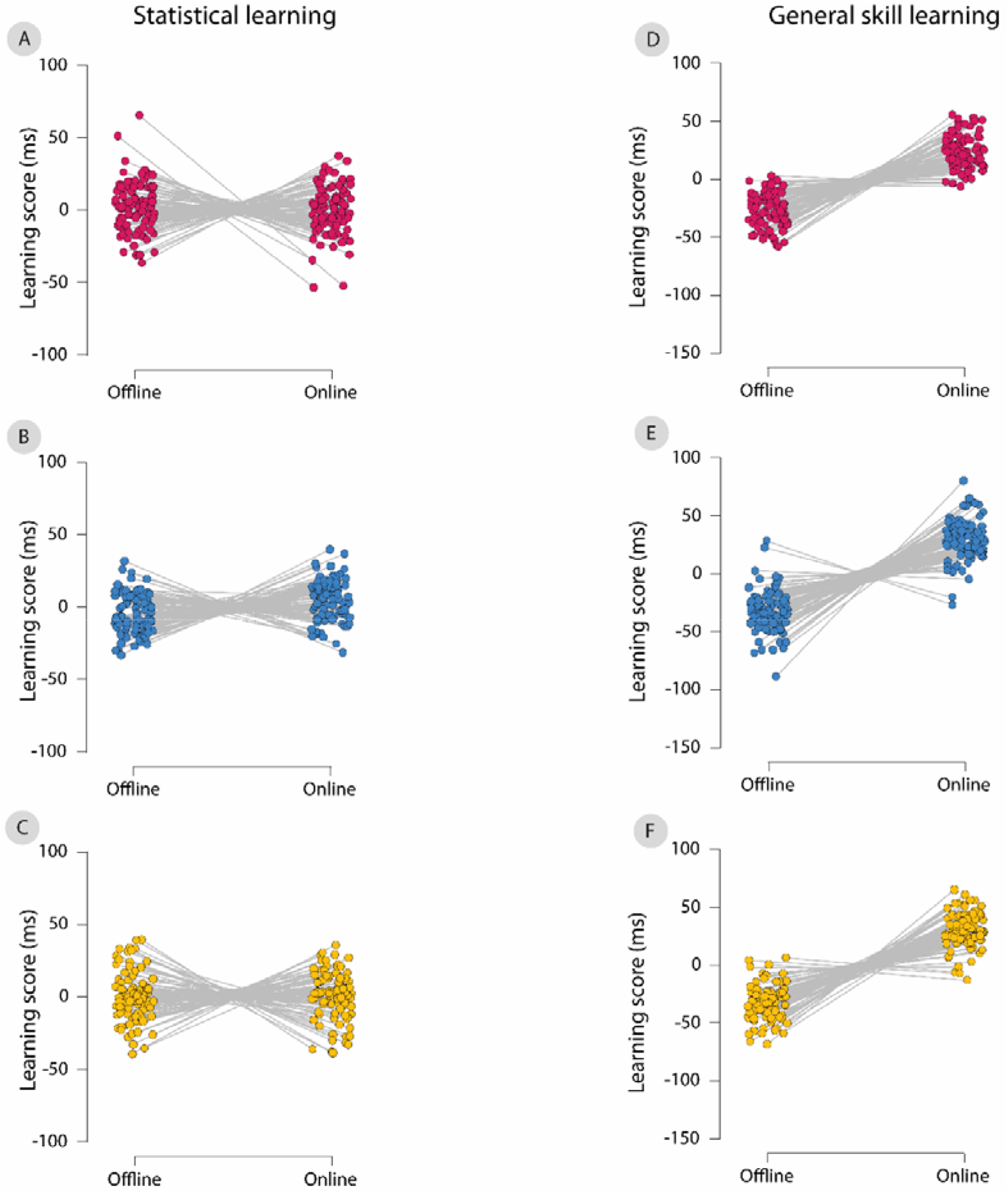
Dynamics of offline/online statistical learning and forgetting in the different groups. The y-axes indicate offline and online learning in milliseconds; the x-axes show the mean offline/online learning score of each participant. The different figures depict the individual data of offline and online statistical learning scores in (A) the self-paced group, (B) the 15-sec group, (C) the 30-sec group, and the individual data of offline and online general skill learning scores in D) the self-paced group, E) the 15-sec group, F) the 30-sec group.

### How did break duration affect offline and online general skill learning?

A mixed-design ANOVA was run on the change scores of general skill learning, with the within-subjects factor of Learning Phase (offline vs. online) and the between-subject factor of Group (self-paced, 15-sec breaks, 30-sec breaks). We found the main effect of Learning Phase [*F*(2, 265) = 920.49, *p* < .001, η_*p*_^2^ = 0.77, *BF*_exclusion_ < 0.001], with a slowing down of RT during the blocks, while an acceleration of RTs occurred after the rests. No main effect of Group was found [*F*(2, 265) = 0.02, *p* = .98, η_*p*_^2^ < .001, *BF*_exclusion_ = 45.61]. The interaction between the Learning Phase and Group factors were significant [*F*(2, 265) = 4.38, *p* = .01, η_*p*_^2^ < .03, *BF*_exclusion_ = 0.01]. However, no differences survived Bonferroni-corrected between-group comparisons for online and offline changes (all comparisons between groups revealed *p* > .17) (Figure 3).

One-sample t-tests revealed that participants learned online (*t*(267) = 29.14, *p* < .001) and forgot offline (*t*(267) = −30.60, *p* < .001) the general skill in the whole sample. This pattern was observed in all three groups. Online learning of the general skill took place in the self-paced group (*t*(87) = 15.87, *p* < .001), in the 15-sec group *t*(89) = 16.35, *p* < .001), as well as in the 30-sec group *t*(89) = 19.18, *p* < .001). In the offline periods, participants’ general skill performance decreased in the self-paced group (*t*(87) = −17.16, *p* < .001), in the 15-sec group (*t*(89) = −16.78, *p* < .001, as well as in the 30-sec group *t*(89) = −20.21, *p* < .001) (see Figure 4D, Figure 4E and Figure 4F).

## Discussion

Our study aimed at testing whether the duration of short rest periods, when neural replay occurs, influences statistical learning and general skill learning. To measure independently these two aspects of learning, we used an implicit sequence learning task, the ASRT task (Howard et al., 2004). We varied the lengths of rest periods across participants: 15 seconds (15-sec group) or 30 seconds (30-sec group) between the learning blocks or participants could decide when to resume the task (self-paced group). We wondered (1) whether the three groups differed in the extent of general skill learning and statistical learning and (2) whether rapid consolidation emerged during between-block rest periods in general skill learning and statistical learning. Break duration affected differently general skills and statistical learning. We observed that the self-paced group was generally slower than the other two groups. However, all groups showed a similar degree of statistical learning. Due to the same proportion of those who learned or forgot offline/online, group level offline and online learning could not be detected in the self-paced and 30-sec groups, while the 15-sec group showed mainly online improvement and offline forgetting (see details in the, Dynamics of offline/online statistical learning and forgetting in the different groups” section of the Supplementary Materials).

Our results suggest that the duration of rest periods is not necessarily decisive in statistical learning over the entire task. This result seems to be inconsistent with the results of Bönstrup and colleagues (2019). They showed that short, 10-second rest periods could facilitate motor skill learning, and this improvement could continue with even shorter rest periods (Bönstrup et al., 2020). In contrast, previous studies that also measured pure statistical learning are consistent with our results (Fanuel et al., 2022). The task used in the study of Bönstrup and colleagues (2020) does not allow the differentiation of subprocesses of learning and mixes general skill learning with statistical learning; therefore, it is difficult to decide which was the determining factor in this result.

According to the results of Buch and colleagues (2021), we expected that the longer rest period (i.e., 30-sec) will result in better learning compared to the shorter rest period (i.e., 15-sec), because it may contain more replays, which is the neural basis of rapid consolidation. However, we measured only one learning session, and it is conceivable that the beneficial effect of expanded rest period would appear rather in a longer run. On the other hand, the length of rest periods used in our study might not have been suitable to capture the critical period when rapid consolidation is beneficial in statistical learning. These questions should be further explored using a much more comprehensive range of rest periods and introducing delayed testing of implicit statistical learning.

In general skill learning, participants showed longer RTs in the self-paced condition where they were allowed to decide about the rest period duration, compared to those conditions where rest period duration was fixed (i.e., 15-sec and 30-sec). How could we interpret the longer RTs in the self-paced group? On the one hand, this difference could be due to a difference in the rest period duration in the self-paced group compared to the two fixed rest period groups. On the other hand, it could be due to the specificity of the self-paced condition. The mean rest period in the self-paced group was similar to the rest period duration of the 15-sec group but the two groups still significantly differed in overall speed. However, the high standard deviation shows considerable variability in the time the participants decided to rest between blocks. We suggest that it is not the duration of the rest period that is critical in the performance of general skill learning, but the nature of the expiry of the rest period (voluntary or compulsory). The knowledge that the rest period will be limited might have urged participants to complete the task as soon as possible, which resulted in faster RTs.

Our results about the offline and online changes in general skill learning are in accordance with previous studies with the same sequence learning task (Fanuel et al., 2022; Quentin et al., 2021): during practice, the speed decreases, and between blocks, it increases. However, our results only partially replicated previous findings on statistical learning. Previously, it was shown that statistical learning mainly occurs during blocks and forgetting between blocks. In our study, this pattern was only detectable for the 15-sec group: in the 30-sec and self-paced groups, such strong dissociation could not be seen in the online vs. offline changes. It is possible that the 30-sec and the self-paced group took enough break time to benefit from both online learning and rapid consolidation (potentially allowing more replay to occur in the offline periods), but the fixed 15-sec length was not enough for the latter. This hypothesis could be supported by the results of Prashad and colleagues (2021), who found offline improvement in probabilistic sequence learning with two-minute-long breaks between the learning blocks (Prashad et al., 2021). However, as the difference in break durations are relatively large between these two studies, it is still a pending question to establish the minimum length of a between-block-break for rapid consolidation. Studies that directly manipulate the number of neural replays between block periods are warranted.

Another possible explanation might be that our study was completed online: participants completed the task in their environment, where the stress level is possibly smaller than in laboratory settings. The limited rest period could have increased the stress level during the experiments, creating similar circumstances that participants experienced in the laboratory. As statistical learning is affected by stress levels (Tóth-Fáber et al., 2021), this could have prompted participants to maximize their performance during practice and benefit from rests more. However, no difference in learning outcomes was found between the groups, suggesting that different lengths of the rest periods only change the learning dynamics but do not affect the outcomes of statistical learning.

Taken together, we observed that the manipulation of the length of the rest periods – indirectly the neural replay – affects general speed on a sequence learning task. In contrast, statistical learning seems to be independent of the length of the rest period. Although the length of rest periods did not affect the outcome of statistical learning, but the dynamics of learning (whether it occurs online or offline): if the brain does not have enough time during breaks for offline consolidation, it might compensate by increasing our online learning performance. Thus, our results suggest that the length of short rest periods has a different effect on separate learning and consolidation processes. Also, from a methodological perspective, our results show the importance to measure the temporal dynamics of learning and not only provide a general measure of the overall learning across the task.

## Acknowledgment

This project was supported by a Gorilla grant that provided free online task hosting (to L. F.). This research was supported by the IDEXLYON Fellowship of the University of Lyon as part of the Programme Investissements d’Avenir (ANR-16-IDEX-0005) (to D. N.); the ATIP-Avenir program (to R. Q), National Brain Research Program (project 2017–1.2.1-NKP-2017-00002). Project no. 128016 has been implemented with the support provided by the Ministry of Innovation and Technology of Hungary from the National Research, Development and Innovation Fund, financed under the NKFI/OTKA K funding scheme (to D. N.). Prepared with the support of the Richter Gedeon Talentum Foundation established by Richter Gedeon Plc. (headquarters: 1103 Budapest, Gyömrői út 19-21.), in the framework of the “Richter Gedeon Excellence PhD Scholarship” (to L. Sz-B.).

## References

Anwyl-Irvine, A., Dalmaijer, E.S., Hodges, N., Evershed, J.K., 2021. Realistic precision and accuracy of online experiment platforms, web browsers, and devices. Behav. Res. Methods. https://doi.org/10.3758/s13428-020-01501-5

Anwyl-Irvine, A.L., Massonnié, J., Flitton, A., Kirkham, N., Evershed, J.K., 2020. Gorilla in our midst: An online behavioral experiment builder. Behav. Res. Methods. https://doi.org/10.3758/s13428-019-01237-x

Barnes, K.A., Howard, J.H., Howard, D. V., Gilotty, L., Kenworthy, L., Gaillard, W.D., Vaidya, C.J., 2008. Intact Implicit Learning of Spatial Context and Temporal Sequences in Childhood Autism Spectrum Disorder. Neuropsychology. https://doi.org/10.1037/0894-4105.22.5.563

Bennett, I.J., Howard, J.H., Howard, D. V., 2007. Age-related differences in implicit learning of subtle third-order sequential structure. Journals Gerontol. - Ser. B Psychol. Sci. Soc. Sci. https://doi.org/10.1093/geronb/62.2.P98

Bönstrup, M., Iturrate, I., Hebart, M.N., Censor, N., Cohen, L.G., 2020. Mechanisms of offline motor learning at a microscale of seconds in large-scale crowdsourced data. npj Sci. Learn. https://doi.org/10.1038/s41539-020-0066-9

Bönstrup, M., Iturrate, I., Thompson, R., Cruciani, G., Censor, N., Cohen, L.G., 2019. A Rapid Form of Offline Consolidation in Skill Learning. Curr. Biol. https://doi.org/10.1016/j.cub.2019.02.049

Buch, E.R., Claudino, L., Quentin, R., Bönstrup, M., Cohen, L.G., 2021. Consolidation of human skill linked to waking hippocampo-neocortical replay. Cell Rep. https://doi.org/10.1016/j.celrep.2021.109193

Cleeremans, A., Jimenez, L., 1998. Implicit sequence learning: The truth is in the details, in: Handbook of Implicit Learning.

de Leeuw, J.R., 2015. jsPsych: A JavaScript library for creating behavioral experiments in a Web browser. Behav. Res. Methods. https://doi.org/10.3758/s13428-014-0458-y

Du, Y., Prashad, S., Schoenbrun, L., Clark, J.E., 2016. Probabilistic motor sequence yields greater offline and less online learning than fixed sequence. Front. Hum. Neurosci. https://doi.org/10.3389/fnhum.2016.00087

Fanuel, L., Pleche, C., Vékony, T., Janacsek, K., Nemeth, D., Quentin, R., 2022. How does the length of short rest periods affect implicit probabilistic learning? Neuroimage: Reports 2, 100078. https://doi.org/10.1016/J.YNIRP.2022.100078

Howard, D. V., Howard, J.H., Japikse, K., DiYanni, C., Thompson, A., Somberg, R., 2004. Implicit Sequence Learning: Effects of Level of Structure, Adult Age, and Extended Practice. Psychol. Aging. https://doi.org/10.1037/0882-7974.19.1.79

Jacoby, L.L., 1991. A process dissociation framework: Separating automatic from intentional uses of memory. J. Mem. Lang. https://doi.org/10.1016/0749-596X(91)90025-F

Janacsek, K., Fiser, J., Nemeth, D., 2012. The best time to acquire new skills: Age-related differences in implicit sequence learning across the human lifespan. Dev. Sci. https://doi.org/10.1111/j.1467-7687.2012.01150.x

Kirchner, W.K., 1958. Age differences in short-term retention of rapidly changing information. J. Exp. Psychol. https://doi.org/10.1037/h0043688

Nemeth, D., Janacsek, K., Londe, Z., Ullman, M.T., Howard, D. V., Howard, J.H., 2010. Sleep has no critical role in implicit motor sequence learning in young and old adults. Exp. Brain Res. https://doi.org/10.1007/s00221-009-2024-x

Prashad, S., Du, Y., Clark, J.E., 2021. Sequence structure has a differential effect on underlying motor learning processes. J. Mot. Learn. Dev. https://doi.org/10.1123/JMLD.2020-0031

Quentin, R., Fanuel, L., Kiss, M., Vernet, M., Vékony, T., Janacsek, K., Cohen, L.G., Nemeth, D., 2021. Statistical learning occurs during practice while high-order rule learning during rest period. npj Sci. Learn. https://doi.org/10.1038/s41539-021-00093-9

Robertson, E.M., Pascual-Leone, A., Miall, R.C., 2004a. Current concepts in procedural consolidation. Nat. Rev. Neurosci. https://doi.org/10.1038/nrn1426

Robertson, E.M., Pascual-Leone, A., Press, D.Z., 2004b. Awareness Modifies the Skill-Learning Benefits of Sleep. Curr. Biol. https://doi.org/10.1016/S0960-9822(04)00039-9

Tóth-Fáber, E., Janacsek, K., Szőllősi, Á., Kéri, S., Nemeth, D., 2021. Regularity detection under stress: Faster extraction of probability-based regularities. PLoS One. https://doi.org/10.1371/journal.pone.0253123

Vékony, T., 2021. ASRT Process Dissociation Procedures (PDP) Task created with jsPsych (Version 1.0.0) [Computer software]. https://doi.org/10.5281/zenodo.7253644

Vékony, T., 2021. Alternating Serial Reaction Time Task created with jsPsych (Version 1.0.0) [Computer software]. https://doi.org/10.5281/zenodo.7124730

Vékony, T., 2021. Verbal N-back task created with jsPsych (Version v1.0.0) [Computer software] https://doi.org/10.5281/zenodo.7100178

Walker, M.P., Brakefield, T., Morgan, A., Hobson, J.A., Stickgold, R., 2002. Practice with sleep makes perfect: Sleep-dependent motor skill learning. Neuron. https://doi.org/10.1016/S0896-6273(02)00746-8

